# Parallel pattern of differentiation at a genomic island shared between clinal and mosaic hybrid zones in a complex of cryptic seahorse lineages

**DOI:** 10.1101/161786

**Authors:** Florentine Riquet, Cathy Liautard-Haag, Lucy Woodall, Carmen Bouza, Patrick Louisy, Bojan Hamer, Francisco Otero-Ferrer, Philippe Aublanc, Vickie Béduneau, Olivier Briard, Tahani El Ayari, Sandra Hochscheid, Khalid Belkhir, Sophie Arnaud-Haond, Pierre-Alexandre Gagnaire, Nicolas Bierne

**Affiliations:** Institut des Sciences de l’Evolution de Montpellier, Université Montpellier, Montpellier, France; CNRS Institut des Sciences de l’Evolution, UMR5554 UM-CNRS-IRD-EPHE, Station Marine OREME, Sète, France; Department of Zoology, University of Oxford, John Krebs Field Station, Wytham, OX2 8QJ, UK; Natural History Museum, Cromwell Road, London SW7 5BD, UK; Department of Genetics, Faculty of Veterinary Science, Universidade de Santiago de Compostela, Campus de Lugo, Lugo, Spain; University of Nice Sophia Antipolis, ECOMERS Laboratory, Faculty of Sciences, Parc Valrose, Nice, France; Association Peau-Bleue, 46 rue des Escais, Agde, France; Center for Marine Research, Ruder Boskovic Institute, Giordano Paliaga 5, 52210 Rovinj, Croatia; Grupo en Biodiversidad y Conservación, IU-ECOAQUA, Universidad de Las Palmas de Gran Canaria, Crta. Taliarte s/n, 35214 Telde, Spain; Institut océanographique Paul Ricard, Ile des Embiez, Six-Fours-les-Plages, France; Océarium du Croisic, Avenue de Saint Goustan, Le Croisic, France; Aquarium de Biarritz, Biarritz Océan, Plateau de l’Atalaye, Biarritz, France; Stazione Zoologica Anton Dohrn, Department Research Infrastructures for Marine Biological Resources, Aquarium Unit, Napoli, Italy; Ifremer - MARine Biodiversity, Exploitation and Conservation, UMR 9190 IRD-IFREMER-UM-CNRS, Sète, France

**Keywords:** clinal hybrid zone, mosaic hybrid zone, reproductive isolation, local adaptation, ecological speciation, parallel evolution

## Abstract

Diverging semi-isolated lineages either meet in narrow clinal hybrid zones, or have a mosaic distribution associated with environmental variation. Intrinsic reproductive isolation is often emphasized in the former and local adaptation in the latter, although both can contribute to isolation. Rarely these two patterns of spatial distribution are reported in the same study system. Here we report that the long-snouted seahorse *Hippocampus guttulatus* is subdivided into discrete panmictic entities by both types of hybrid zones. Along the European Atlantic coasts, a northern and a southern lineage meet in the southwest of France where they coexist in sympatry with little hybridization. In the Mediterranean Sea, two lineages have a mosaic distribution, associated with lagoon-like and marine habitats. A fifth lineage was identified in the Black Sea. Genetic homogeneity over large spatial scales contrasts with isolation maintained in sympatry or close parapatry at a fine scale. A high variation in locus-specific introgression rates provides additional evidence that partial reproductive isolation must be maintaining the divergence. Surprisingly, fixed differences between lagoon and marine populations in the Mediterranean Sea belong to the most differentiated SNPs between the two Atlantic lineages, against the genome-wide pattern of structure. These parallel outlier SNPs cluster on a single chromosome-wide island of differentiation. Since Atlantic lineages do not match the lagoon-sea habitat variation, genetic parallelism at the genomic island suggests a shared genetic barrier contributes to reproductive isolation in contrasting contexts -*i.e.* spatial *vs*. ecological. We discuss how a genomic hotspot of parallel differentiation could have evolved and become associated either with space or with a patchy environment in a single study system.

## Introduction

The spatial context of contact zones between partially isolated taxa and their relationship with environmental variation was long thought to offer great promises to unravel the nature and origin of species. Though each taxon may be genetically homogeneous over large distances, they often meet in abrupt genetic discontinuities, called hybrid zones, in which partial reproductive isolation limits gene exchange (Barton and Hewitt 1985, Hewitt 1988). Hybrid zones are extremely informative for exploring the genetic basis of reproductive isolation (e.g. Teeter et al. 2008, Christe et al. 2016) and local adaptation (e.g. Jones et al. 2012, Larson et al. 2013, Soria-Carrasco et al. 2014), as well as identifying genomic regions involved either in increased genomic differentiation (Ravinet et al. 2017) or adaptive introgression (Hedrick 2013). The hybrid zone literature usually contrasts two spatial patterns (Harrison 1993): (i) clinal hybrid zones, with parapatrically distributed parental forms on both sides of a genetic divide, and (ii) mosaic hybrid zones, when the environment consists of a mosaic of habitat patches to which taxa (ecotypes, host races, hybridizing species) are somehow specialized. Contrasting with this long-standing dichotomy, hybrid zones of both types have now been recognized to be multifactorial and maintained by exogenous and endogenous diverging mechanisms (*i.e.* local adaptation and intrinsic reproductive isolation, respectively; Barton and Hewitt 1985, Bierne et al. 2011). Nonetheless, clinal hybrid zones still tend to be interpreted as being mainly maintained by intrinsic reproductive isolation evolved in allopatry before contact (the tension zone model, Barton and Hewitt 1985). Conversely, local adaptation is often emphasized to drive mosaic distributions which are interpreted as evidence that selection at local adaptation genes is acting to increase differentiation between habitats in a parallel fashion (Nosil and Feder 2012). This sketchy dichotomy is anchored by emblematic study systems for which decades of research allow support for such interpretations. For instance, the hybrid zone between the mice *Mus musculus musculus* and *M. m. domesticus* in central Europe (Boursot et al. 1993) has been well demonstrated to be maintained by selection against hybrid genotypes (Britton-Davidian et al. 2005, Good et al. 2008) after secondary contact (Duvaux et al. 2011). Although the position of the hybrid zone was initially found to be associated with rainfall (Hunt and Selander 1973), local adaptation is not considered to contribute much to the isolation. At the other extreme, local adaptation of the same genetic variants has repeatedly allowed marine three-spined sticklebacks to evolve into a freshwater ecotype (Jones et al. 2012). Intrinsic selection is thought absent in the marine-freshwater sticklebacks system, although hybrids perform poorly in both environment and tend to reside in salinity ecotones (Vines et al. 2016). It would be misleading, however, to suggest the two alternative spatial contexts and relations to environmental variation may correspond to alternative routes to speciation (e.g. mutation-order *vs.* ecological speciation). The list of hybrid zones maintained both by local adaptation and intrinsic reproductive isolation is also long. *Bombina* toads (Szymura and Barton 1986), *Gryllus* crickets (Rand and Harrison 1989, Larson et al. 2014), or *Mytilus* mussels (Bierne et al. 2003) are well-known examples of mosaic hybrid zones maintained by exogenous and endogenous selection. However, we do not usually expect intrinsic incompatibilities to be reused in a parallel fashion. Besides, parallel genetic divergence associated with contrasting environmental conditions (e.g. marine/freshwater, highland/lowland, host races) remains a strong hallmark of ecologically-driven divergence (Bierne et al. 2013). Given this -hybrid zone-context, in this paper we aim to provide an example of a genetic parallelism with a lack of apparent ecological convergence. We expect this counter-example could contribute to break the inductive reasoning that genetic parallelism means ecological convergence. However, as we discovered this pattern by serendipity in a newly studied complex of cryptic genetic backgrounds in a non-model system, we have to describe this system first.

We studied the population genetics of the long-snouted seahorse, *Hippocampus guttulatus,* across a large part of its geographic range. We developed an assay of 286 informative SNPs chosen from more than 2,500 SNPs identified in a population transcriptomic study (Romiguier et al. 2014). *H. guttulatus* displays poor dispersal abilities (e.g. site-fidelity, weak swimming performance, lack of dispersive stage) and inhabits fragmented coastal habitats along its distribution range (from the English Channel through the Mediterranean and Black Seas, Lourie and Vincent 2004). In addition most populations are small and patchy. Given these biological characteristics, a strong genetic structure could have been expected. In agreement, a very low genetic diversity was observed in *H. guttulatus* when compared to 75 non-model animal species (Romiguier et al. 2014). However, genetic differentiation proved to be very weak over very large geographic distances based on the two genetic studies conducted to date with microsatellite loci (e.g. from the United Kingdom to North of Spain, Woodall et al. 2015, or across the Cape Finisterre oceanographic barrier, López et al. 2015). Four well-differentiated genetic clusters, each distributed over extended regions, were delineated by genetic discontinuities corresponding to usual delimitations between vicariant marine lineages (Woodall et al. 2015) – between the Iberian Peninsula and the Bay of Biscay in the North Eastern Atlantic, between the Atlantic Ocean and the Mediterranean Sea, and between the Mediterranean and Black Seas. Although such genetic differentiation matches well with oceanographic barriers and was interpreted as spatial differentiation, this pattern is also concordant with the existence of reproductively isolated cryptic lineages, which boundaries were trapped by exogenous barriers. In this latter interpretation, although the location of genetic breaks would be due to exogenous factors (e.g. temperature, salinity or oceanic fronts), the barrier to gene flow would mainly be driven by barrier loci that restrict gene flow on a large fraction of the genome (*i.e.* the coupling hypothesis; Bierne et al. 2011, Gagnaire et al. 2015, Ravinet et al. 2017). This hypothesis is receiving increasing support (e.g. Le Moan et al. 2016, Rougeux et al. 2016, Rougemont et al. 2016) and could well explain the genetic structure observed in the long-snouted seahorse.

Using newly developed SNP-markers spread along the genome and a more extensive sampling along the *H. guttulatus* distribution range, we challenged the initial interpretation of barriers to dispersal against the alternative hypothesis of reproductive isolation between semi-isolated genetic backgrounds. We described five cryptic semi-isolated lineages: two lineages in the Atlantic Ocean with a parapatric distribution, two lineages in the Mediterranean Sea with a patchy fine-grained environment association (lagoon *vs.* marine environments), and one in the Black Sea. Surprisingly, a shared genomic island of clustered outlier loci was involved both in the isolation between the two parapatric lineages in the Atlantic Ocean and between the marine and lagoon ecotypes in the Mediterranean Sea. Furthermore, the North Atlantic lineage was related to the lagoon ecotype at this genomic island, against the genome-wide pattern of structure. However, the two Atlantic lineages inhabit both marine and lagoon habitats. We argue that the *H. guttulatus* complex could become one of a few systems where a clinal and a mosaic hybrid zone are observed concomitantly, and a valuable new counter-example that provides evidence of genetic parallelism in absence of ecological convergence.

## Materials and Methods

### Sampling and DNA extraction

*H. guttulatus* samples were collected alive from 25 sites (Fig. 1) using a variety of methods (snorkeling, scuba diving, trawling nets, aquarium, donations). The dorsal fin of each individual was clipped using a non-lethal procedure (Woodall et al. 2012), before releasing back the individual into its natural habitat. In three sites (sites 14, 18 et 19 in Fig. 1), dorsal fins were clipped from captive-bred seahorse held at the Mare Nostrum Aquarium, France, recently sampled in their natural habitats. Each individual sample was preserved and stored in 96% ethanol for subsequent genetic analyses.

**Figure 1.**
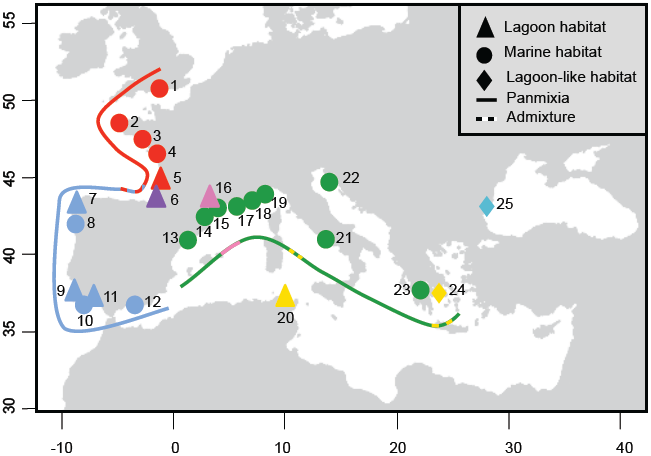
Sampling locations of *Hippocampus guttulatus.* Each study site is labeled as follow: 1-Poole, United Kingdom, 2-Brest, France, 3-Le Croisic, France, 4-Ré Island, France, 5-Arcachon, France, 6-Hossegor, France, 7-Coruna, Spain, 8-Vigo, Spain, 9-Portimao, Portugal, 10-Faro (maritime site), Portugal, 11-Faro (lagoon site), Portugal, 12-Malaga, Spain, 13-Tossa de Mar, Spain, 14-Leucate, France, 15-Sète (maritime site), France, 16-Thau lagoon, France, 17-La Ciotat, France, 18-Le Brusc, France, 19-Cavalaire-sur-Mer, France, 20-Bizerte lagoon, Tunisia, 21-Naples, Italy, 22-Rovinj, Croatia, 23-Kalamaki, Greece, 24-Halkida, Greece, and 25-Varna, Bulgaria. Lagoon habitats are represented by triangles, lagoon-like habitats by diamonds and maritime habitats by circles. Red, blue, green, pink and turquoise symbols stand respectively for the North Atlantic, South Atlantic, Mediterranean Sea, Mediterranean lagoon, and Varna cluster, all of them showing panmixia (solid lines). Dashed lines stand for contact zones (sites 6, 20 and 23).

Whole genomic DNA was extracted following either Woodall et al. (2015), López et al. (2015) or using a standard CetylTrimethyl Ammonium Bromide (CTAB) Chloroform:Isoamyl alcohol (24:1) protocol (Doyle and Doyle 1987). For samples from Vigo (Spain), the whole genome was amplified using the GenomiPhi kit (GE HealthCare) according to the manufacturer’s protocol. Quality and quantity of DNA extraction was checked on an agarose gel, and normalized to 35 ng.µL^−1^ using Qubit Fluorometric Quantitation (Invitrogen).

### Data mining for SNP markers

A set of 12,613 contigs was examined to identify SNPs. This included one mitochondrial contig (GenBank accession number: AF192664) and 12,612 contigs from Romiguier et al. (2014). Briefly, Romiguier et al. (2014) produced high-coverage transcriptomic data (RNAseq) for six *H. guttulatus* from three locations (Le Croisic, Atlantic Ocean, France; Faro, Atlantic Ocean, Portugal; Thau lagoon, Mediterranean Sea, France), and two *H. hippocampus* from two locations (Sète, Mediterranean Sea, France; Bizerte, Mediterranean Sea, Tunisia). *De novo* transcriptome assembly based on Illumina reads was performed using a combination of the programs ABySS (Simpson et al. 2009) and Cap3 (Huang and Madan 1999), then mapped to predicted cDNAs (contigs) with BWA (Li and Durbin 2009). Based on these 12,613 contigs, SNPs were identified using the bioinformatic pipeline described in Bouchemousse et al. (2016). SNPs were called with Read2SNPs (Gayral et al. 2013) and filtered out according to the following criteria to exclude: 1) SNPs showing more than two alleles, 2) SNPs failing to be sequenced in at least one location, 3) SNPs present in only one individual (*i.e.* singletons), 4) SNPs identified as paralogs using the paraclean option of Read2SNPs, and 5) SNPs closer than 20 bp from a contig extremity or an exon limit when blasted against the stickleback, cod and tilapia genomes. This resulted in 2,684 selected SNPs screened with the Illumina Assay Design Tool (ADT) software to select 384 SNPs on the basis of their quality index (ADT score > 0.6). An Illumina BeadXpress^®^ with Veracode^TM^ technology (GoldenGate^®^ Genotyping Assay) was then used to genotype the 384 selected SNPs.

To identify their chromosomal positions, the template sequences of the targeted SNP were blasted against i) the genome of the Gulf pipefish (*Syngnathus scovelli*, Small et al. 2016) and ii) the scaffolds of the tiger tail seahorse (*Hippocampus comes*, Lin et al. 2016). *H. comes* scaffolds being unplaced, we blasted these scaffolds against *S. scovelli* genome and, for more consistency, blatted (Bhagwatt et al. 2012) them against seven well-assembled fish genomes; zebrafish (*Dano rerio*, Howe et al. 2013), fugu (*Takifugu rubripes*, Kai et al. 2011), tetraodon (*Tetraodon nigroviridis*, Jaillon et al. 2004), Nile tilapia (*Oreochromis niloticus*, Brawand et al. 2014), medaka (*Oryzias latipes*, Kasahara et al. 2007), stickleback (*Gasterosteus aculeatus*, Jones et al. 2012), and European seabass (*Dicentrarchus labrax*, Tine et al. 2014).

SNPs were polarized using *H. hippocampus* as an outgroup to identify the most parsimonious ancestral variant, which allowed the derived allele state to be identified. The Joint Site-Frequency Spectrum (JSFS is the more informative summary statistics regarding inter-population polymorphism; Wakeley 2008) obtained from the original transcriptome-wide SNP dataset (Romiguier et al. 2014) was compared to the JSFS obtained by the subset of 384 SNPs to investigate the extend of ascertainment bias potentially induced by our marker selection. In order to compare the JSFS obtained with both datasets, the JSFS dimension was projected down to a 5×5 matrix, which was the dimension of the Romiguier et al. (2014) dataset.

### Analyses of Genetic Diversity and Genetic Structure

The mean number of alleles per locus (*N*_*all*_) and the allelic richness (*A*_*r*_, *i.e.* the expected number of alleles corrected for sampling size based on a rarefaction method) were estimated using the function divBasic implemented in the R package diveRsity (Keenan et al. 2013). Allelic frequencies, expected heterozygosity (*H*_*e*_) and fixation index (*F*_*IS*_) were estimated using GENEPOP on the web (Raymond and Rousset 1995, Rousset 2008). To get a genome-wide picture from the multi-locus genotype dataset summarizing inter-population polymorphism, the raw SNP data were visualized by INTROGRESS (Gompert and Buerkle 2010), although its primary goal is to estimate genomic clines, but allows in a simple manner to plot raw data to investigate any genomic signal. Alleles derived from each of the parental populations were counted at the individual level and converted into a matrix of counts, then used to visualize the multilocus genotype of each individual.

Genetic structure among sampling sites was depicted using both Principal Component Analysis (PCA) computed on the matrix of genotypes, and an individual-based Bayesian clustering method. The latter being a model-based approach with strong priors and hypotheses (Hardy-Weinberg equilibrium, no linkage disequilibrium) contrasted with PCA, a distant-based approach when few (nearly no) assumptions may be violated. Comparing results from both analyses using different statistical approaches, allows us to make solid assumptions about our data. The PCAs were carried out using the R package ADEGENET 1.4–2 (Jombart 2008, Jombart et al. 2011). The individual-based Bayesian clustering analysis was performed with the software STRUCTURE 2.3.4 (Pritchard et al. 2000, Falush et al. 2003). For each value of *K* (ranging from 1 to 25), 30 replicate chains of 150,000 Monte Carlo Markov Chain (MCMC) iterations were run after discarding 50,000 burn-in iterations. An admixture model with correlated allele frequencies was applied with *a priori* information on samples origin. Note that this method makes the assumption of homogeneous admixture rate in the genome (neutral admixture) and therefore return a sort of weighted average admixture rate when introgression is heterogeneous across the genome. To determine individual ancestry proportions (*q*-values) that best matched across all replicate runs, CLUMPP (Jakobsson and Rosenberg 2007) was used and individuals’ assignment visualized in the R software.

Once the different genetic clusters were identified (and cross-validated using both methods), genetic homogeneity among samples belonging to a cluster was checked, allowing subsequently pooling of samples into clusters with a minimum of seven individuals per cluster. Genetic structure was computed among clusters by calculating global and pairwise *F*_*ST*_ (Weir and Cockerham 1984) using GENEPOP on the web (Rousset 2008). Exact tests for population differentiation (10,000 dememorization steps, 500 batches and 5,000 iterations per batch) were carried out to test for differences in allele frequencies. s, defined as the adjusted *p*-values using an optimized False Discovery Rate approach, were computed using the QVALUE package in the R software (Storey 2002) to correct for multiple testing.

Evolutionary history of genetic clusters was also investigated under a model of divergence and admixture events using the population graph approach implemented in the TREEMIX software (Pickrell and Pritchard 2012). This software uses the covariance matrix of allele frequency between pairs of populations to infer both population splits and gene flow. A maximum likelihood population tree is first generated under the hypothesis of an absence of migration, and admixture events are sequentially added, improving (or not) the tree model. This statistical method shows the benefit of constructing population trees while testing for gene flow between diverged populations at the same time. Samples with too small sample size (N<7) or without random mating (see Results) were removed as they generate erroneous results (Pickrell and Pritchard 2012). Using the total data set, *i.e.* 286 loci, five migration events were sequentially added to look for the best tree to fit the data and the number of migration events that reached an asymptotic likelihood was kept.

### Outlier detection

The use of several independent methods is often recommended to improve accuracy of outlier loci detection (Pérez-Figueroa et al. 2010, de Villemereuil et al. 2014). Each method is differently impacted by the genetic structure and/or demographic history of the study species, which is usually unknown, leading to frequent inconsistencies across methods (Gagnaire et al. 2015). To cope with these problems, outlier loci were detected using four different methods. First, BAYESCAN (Foll and Gaggiotti 2008) is a Bayesian method that uses a logistic regression model to estimate directly the posterior probability that a given locus is under selection. We used default parameter values in our analyses to detect outliers among the clusters previously identified. The second approach (Duforet-Frebourg et al. 2014) implemented in the R package PCAdapt (Luu et al. 2016) is based on a hierarchical model where population structure is first depicted using K factors. No *a priori* on the genetic structure (and thus, no clustering) is required in advance. Loci that are atypically related to population structure, measured by the K factors, are identified as outliers. For each value of K (ranging from one to ten), ten replicate chains of 150,000 Monte Carlo Markov Chain iterations were performed and we discarded the first 5,000 iterations as burn-in. The third approach uses the estimated co-ancestry matrix to compute an extension of the original Lewontin and Krakaeur statistic that account for the history of populations under a model of pure drift (Bonhomme et al. 2010). Finally we used the custom simulation test described in Fraïsse et al. (2014). The idea of this test is to use simulations of the best-supported demographic model to obtain the neutral envelope of the joint distribution of pairwise F_ST_ in a four-population analysis. This test uses the fact that it is easy to have false positives in each of two pairwise comparisons but that outliers in both comparisons, against the genome-wide structure, are more likely to be true positives. Roux et al. (2016) found that an Isolation-with-Migration (IM) model fitted the seahorse data well and that more parameterized models did not improve the fit. These authors also found that the time of divergence (Tsplit) was very similar in each pairwise comparison. We therefore used a four-population IM model with the population sizes and migration rates inferred by Roux et al. (2016) on the transcriptome data of Romiguier et al. (2014). The two latter tests are very similar in their spirit, although one use a simple history of divergence with drift and the other intend to explore more complex demographic histories. They were here used to identify parallel SNPs that provide a grouping of populations that goes against the genome-wide trend.

Genotype discordance among loci in admixed samples (*i.e.* genomic cline analysis) was tested using Barton’s concordance method as described in Macholán et al. (2011). The method fits a quadratic function to the relationship between a single locus hybrid index (0 or 1 if homozygous, 0.5 if heterozygous) and the genome-wide hybrid index. The function parameters measure the deviation from x = y (*i.e.* the expectation of homogeneous genome-wide introgression) as a function of the expected heterozygosity. Instead of testing the deviation from the diagonal, our aim was to compare genomic clines (*i.e.* regression curves) between geographic samples, as discordance was observed in one population and not others.

Finally, we tested for gene ontology (GO) terms enrichment to determine if outlier loci displayed functional enrichment, compared to the full dataset, using Fisher’s Exact Test with Multiple Testing Correction of FDR (Benjamini and Hochberg 1995) implemented in the software Blast2GO (Conesa and Götz 2008).

## Results

### SNPs characterization/calling: an efficient genotyping method in a protected species with low genetic diversity

Of the 384 SNPs, including five SNPs that diagnostically distinguished *H. guttulatus* from *H. hippocampus* (one mitochondrial and four nuclear), a total of 318 SNPs amplified successfully. Of these 318 SNPs, 286 showed a minor allele frequency above 5%. Two *H. hippocampus* initially identified as *H. guttulatus* were removed from the initial dataset, resulting in a final dataset of 292 *H. guttulatus* genotyped for 286 SNPs.

In order to evaluate the extent of ascertainment bias that may be induced by our procedure for selecting markers, we compared the Joint Site-Frequency Spectra (JSFS) obtained with the original dataset of Romiguier et al. (2014) -assumed freed from ascertainment bias-with our 314-SNP dataset. JSFSs are detailed in Supporting Information 1 (Fig. SI1). Briefly, as expected, the data of Romiguier et al. (2014) is principally composed of singletons that represented 35-40% of the SNPs (Fig. SI1A), while this proportion was reduced twofold in our 314-SNP dataset (Fig. SI1B). We efficiently removed rare variants without biasing the frequency spectrum too much as the deficit of singletons was homogeneously compensated by every other cells of the JSFSs. An even representation over the entire allele frequency range was indeed observed based on our dataset. The comparison of the two JSFSs (Fig. SI1C) reveals that very few cells apart from singletons have an excess above 5%, suggesting limited ascertainment bias in our SNP panel, except with rare alleles as expected.

With a limited ascertainment bias, with rare alleles being underrepresented, no missing data, and constraints on our model study (small amount of DNA available with the use of non-lethal fin-clipping sampling techniques), selecting SNPs characterized from a preliminary population transcriptomic survey proved to be a straightforward strategy for genome-wide investigation of the spatial distribution in this species compared to classical genotyping-by-sequencing approaches.

### A strong genetic structure delineating five broadly distributed panmictic genetic clusters

Estimates of the mean number of alleles per locus (*N*_*all*_), allelic richness (*A*_*r*_), expected heterozygosity (*H*_*e*_), and departure from Hardy-Weinberg equilibrium (HWE; *F*_*IS*_), for each study site and each genetic cluster identified, are presented in Supporting Information 2. The gene diversity was similar among populations with no significant departure from HWE observed (with the exception of site 6 which is a hybrid population, see below).

The Principal Component Analysis (PCA) revealed clear differentiation separating five clusters (Fig. 2). Along the first two axes (left panel), explaining 60.1% and 16.2% of the variance, the PCA showed clear differentiation between North Atlantic (sites 1-5, in red), South Atlantic (sites 7-12, in blue), Mediterranean Sea (sites 13-25, lagoon site 16 excluded, in green) and Mediterranean Thau lagoon (site 16, in pink). Hossegor individuals (site 6, in purple) clustered either with the North Atlantic (12 individuals) or with the South Atlantic groups (3 individuals). Mediterranean sites spread out along the third axis (9.9% of the variance explained, right panel), with Bizerte (site 20, in gold) being closer to the Mediterranean Thau lagoon on the one hand, and Varna (Black Sea, site 25, in turquoise) standing out from the Mediterranean group on the other hand.

**Figure 2.**
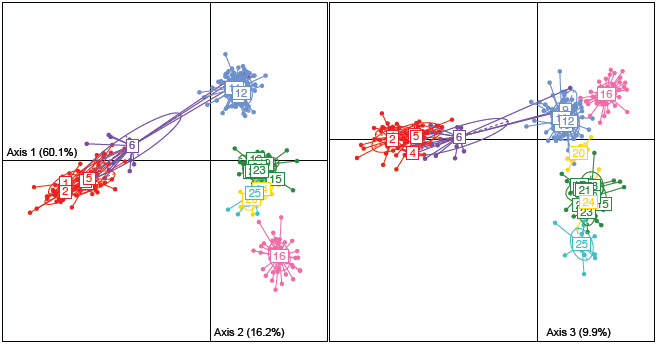
Genetic population structure based on 286 SNP markers analyzed by Principal Component Analyses depicting axis 1 (explaining 60.1% of the variance) and axis 2 (explaining 16.2% of the variance; left panel) and axes 1 and 3 (explaining 9.9% of the variance; right panel). Samples are numbered and colored according to Fig. 1. Each label shows the barycenter of each study site.

This pattern of genetic structure was also detected by the STRUCTURE analysis, of which the output composed of five clusters is presented in Supporting Information S3. By selecting a K-value of five, we distinguished identical clusters composed by: 1-the North Atlantic sites, 2-the South Atlantic sites, 3-the Mediterranean Sea sites, 4-the Mediterranean Thau lagoon and 5-Varna. Similar to the PCA (Fig. 2), a gradient of introgression is visible along the Atlantic coasts, with a decreasing proportion of South Atlantic cluster ancestry North of Hossegor (site 6). While most Hossegor individuals belong to the North Atlantic genetic background, three individuals were genetically assigned to the South Atlantic genetic background. Bizerte (site 20), and to a lesser extent Halkida (site 24), proved to have mixed ancestries from both Mediterranean lagoon and Sea clusters. Increasing the K-value resulted in additional clusters composed by Bizerte (site 20, K=6), Halkida (site 23, K=7), but then (K>7) no individuals were completely assigned to any new cluster.

Altogether, distance-based (PCA) and model-based (Structure) analyses supported the identification of five clusters, a pattern also showed by the visualization of raw mutli-locus genotype data (Supporting Fig. SI3B). This representation illustrates that most markers contribute to the signal of five genetic clusters.

Importantly, no significant departure from panmixia was observed within each cluster (SI2). Furthermore, genetic homogeneity was observed between sites within each cluster (SI2). In contrast, Fisher exact tests revealed significant differences in allelic frequencies among clusters *(p*-value < 0.001), with significant differentiation being observed for all pairwise comparisons (*F*_*ST*_ ranging from 0.024 between the Mediterranean Sea cluster and Halkida to 0.260 between the North Atlantic cluster and Varna).

Finally, the population tree inferred using TREEMIX without accounting for migration (Fig. 3A) was highly consistent with all above analyses. Atlantic clusters branching together on one hand, and Mediterranean lagoons (Thau and Bizerte) branching together on the other hand. Interestingly, three admixture events significantly improved the model as compared to a scenario without migration (*p*-value < 0.001; Fig. 3B). This population tree indicated significant gene flow among four *H. guttulatus* clusters, between the north and the south genetic clusters in the Atlantic coasts, in concordance with the gradient of introgression along the Atlantic coasts (Fig. 2 and SI3), between marine and lagoon samples in the Mediterranean Sea, and finally between the North Atlantic and Mediterranean lagoon samples. We suggest that this latter observation is most probably mainly driven by parallel outlier loci (see below), either as a result of adaptive introgression or as a consequence of a shared history of divergence retained at outlier loci against massive secondary gene flow.

**Figure 3.**
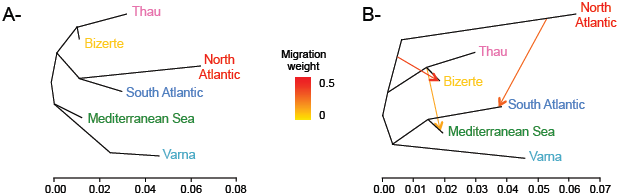
Population trees inferred by TREEMIX (A) without or (B) with 3 migration events. Admixture arrows are colored according to the migration weight. The model including three admixture events significantly improved the fit as compared to a situation without migration (*p*-value < 0.001). Panmictic clusters are colored according to Fig. 1.

### Signature of selection and genetic parallelism

Nine SNPs out of 286 (3.15%) were consistently identified to depart from neutrality with the four tests (BayeScan, PCAdapt, FLK and custom simulation test). Interestingly, six of them showed very similar allele frequency patterns, distinguishing North Atlantic sites, Mediterranean Thau lagoon, Bizerte (site 20) and, in a lesser extent Halkida (site 23) and Varna (site 25) from South Atlantic and Mediterranean Sea sites (Fig. 4), and pointing out genetic parallelism (*i.e.* convergence of allele frequency patterns) between these lineages.

**Figure 4.**
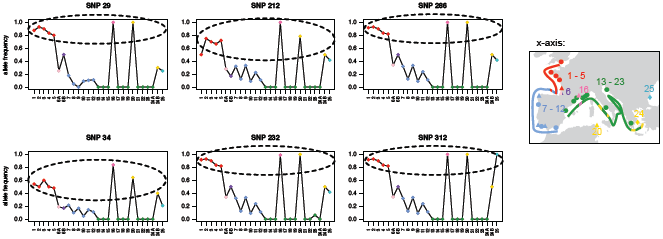
*H. guttulatus* allele frequencies (y-axis) for the six outliers with very similar high allele frequency. Each study site (x-axis) is labeled and colored according to Fig. 1, reminded by a simplified map on the right. Hossegor was separated in 6A and 6B and Halkida in 24A and 24B according to their North or South Atlantic / Mediterranean Thau lagoon or Sea genetic background, respectively (see Fig. 2, SI3).

These six outliers located on different *H. guttulatus* contigs were located on different *Hippocampus comes* scaffolds, except SNPs 29 and 286 mapping to a unique *H. comes* scaffold (Fig. 5A). Interestingly, these scaffolds –that contain outliers-consistently mapped to a unique chromosome in *Syngnathus scovelli* (Fig. 5A). Results were similar when directly blasting these six *H. guttulatus* contigs against *S. scovelli* genome, but with SNP 29 mapping to an unplaced scaffold (Fig. 5A). A unique chromosome is still involved when blatting *H. guttulatus* outlying contigs against seven additional well-assembled fish genomes, in agreement with a well-conserved synteny of fishes (detailed in SI5).

**Figure 5.**
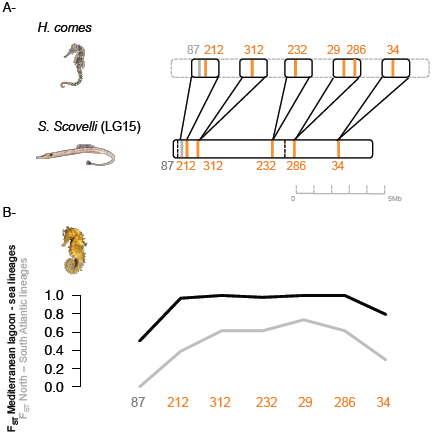
Outlier cross-mapping (A-) and *F*_*ST*_ between the Mediterranean lagoon and Sea lineages (in black), and between the North and South Atlantic lineages (in grey) along the unique chromosome (B-). As *H. comes* scaffolds are unplaced, *H. guttulatus* SNPs were first blasted on *H. comes* scaffolds, then each *H. comes* scaffold was mapped to *S. scovelli* genome. A putative *H. comes* chromosome was hence reconstructed (A-). The order of the scaffolds and SNPs according to the blasts was conserved. Outlier SNPs displaying parallel differentiation between Atlantic lineages and Mediterranean ecotypes are colored in orange, while the grey outlier showed high differentiation between the Mediterranean Thau lagoon and other sites.

Genetic parallelism was visualized by plotting F_ST_ of Mediterranean lagoon and sea locations (x-axis in Fig. 6) against F_ST_ between North and South Atlantic clusters (y-axis in Fig. 6). Outliers showing genetic parallelism appeared in the top right of Figure 6, showing elevated genetic differentiation between Mediterranean lagoon and marine sites (x-axis) on the one hand, and between North and South Atlantic sites (y-axis) on the other hand. This F_ST_ - F_ST_ co-plot also showed how the parallel pattern of differentiation observed at these six loci would be highly unlikely under the inferred demographic history of populations.

**Figure 6.**
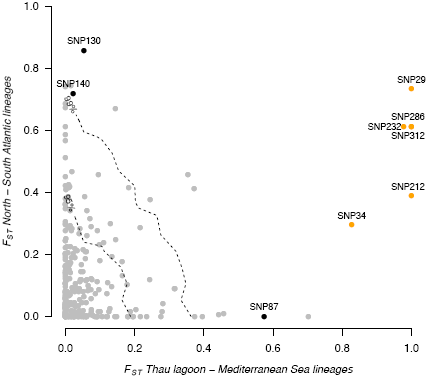
Genome scan of infra-specific differentiation in *H. guttulatus*. Dashed lines represent the 95% and 99% quantiles of the neutral envelope of F_ST_ obtained following Fraïsse et al. (2014). Loci identified by all methods as outliers are colored in orange – the six outliers that displayed parallel differentiation between Atlantic lineages and Mediterranean ecotypes – and in black – the three other outliers.

Three other outliers were also consistently identified (SNPs 87, 130 and 140; Fig. 6 and Supporting Information SI4) using all four methods. They distinguished either the Mediterranean Thau lagoon (*i.e.* high genetic differentiation along the x-axis in Fig. 6; SNP 87), the North Atlantic sites or the South Atlantic sites (*i.e.* high genetic differentiation along the y-axis in Fig. 6; SNPs 130 and 140) from all other sites. By using a similar approach for convergent outliers, we only observed SNP 87 that consistently mapped in the unique chromosome.

Figure 7 shows the genomic cline analysis obtained with SNP 29, an outlier that clustered on the unique chromosome (Fig. 5 and SI5), and SNP 130, an outlier between the northern and the southern Atlantic lineage but that mapped to another chromosome and not differentiated in the Mediterranean Sea (no genetic parallelism). Regressions were found to be different at SNP 29 and other outliers that clustered on the same chromosome, when the Hossegor sample was included in the analysis or not (Fig. 7), while regressions were always close to the diagonal with SNP 130 and other outliers that mapped to other chromosomes. Introgression did not occur at the same rate for the loci mapping to the unique chromosome, with a clear discordance between the clustered outlier loci and the rest of the genome in the Hossegor sample, in which North Atlantic-like seahorses have the South Atlantic allele at the genomic island outliers.

**Figure 7.**
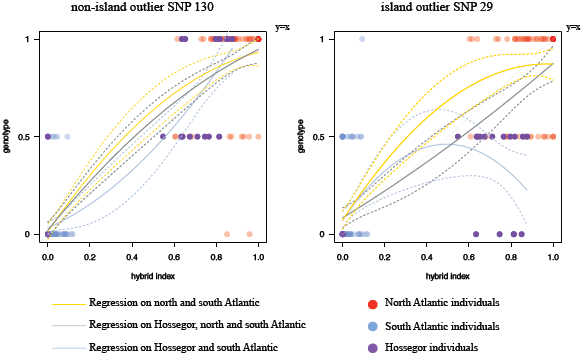
Genomic cline plots for two illustrative markers showing extreme level of differentiation (SNPs 29 and 130). Genomic clines were conducted on North and South Atlantic lineages following polynomial regressions on all Atlantic individuals without Hossegor (in yellow), on all Atlantic individuals (in grey) and on all Atlantic individuals, without North Atlantic (in blue). Dotted lines represent the 95% confidence intervals. Circles indicate the raw genotype data (ancestral homozygotes on the top line –the “1” genotype, heterozygotes in the middle –the “0.5” genotype, and derived homozygotes on the bottom line –the “0” genotype).

Note that of all the 168 fish sexed (out of 292 fishes), comprising individuals from each of the five lineage and contact zones in a balanced sex ratio, loci with extreme level of differentiation could not be related to any sex differences. In addition, the analysis of gene ontology terms for outlier loci did not reveal any significant functional enrichment.

## Discussion

Genetic analyses of the long-snouted seahorse revealed cryptic discrete panmictic genetic lineages that meet either in a narrow contact zone in the Atlantic, or display a mosaic distribution associated with environmental variation in the Mediterranean Sea. Despite limited dispersal abilities and seemingly small population sizes (but see Curtis and Vincent 2006), each lineage showed remarkable genetic homogeneity over very large distances, with genetic panmixia observed within each lineage. This spatial structure, with strong and sharp genetic subdivisions, is not expected if genetic drift was predominantly responsible for the genetic differentiation between these populations.

The spatial organization of the different genetic backgrounds proved to be an unusual combination of geographic subdivision in the Atlantic Ocean, and genetic structure related to the sea-lagoon ecological contrast in the Mediterranean Sea. Surprisingly, these two subdivisions partly relied on the same genetic architecture. We observed genetic parallelism at some markers showing extreme levels of differentiation between habitats in the Mediterranean Sea, but also between geographic lineages in the Atlantic Ocean. Intriguingly, all the loci showing convergent allele frequency patterns mapped to a single chromosome. This tends either to suggest genetic differences do not accumulate evenly across the genome, or that these genomic regions are easier to characterize with low-density genome scans.

We suggest the existence of a shared evolutionary history between Atlantic parapatric lineages and Mediterranean ecotypes, with the Mediterranean lagoon ecotype anciently related to the North-Atlantic lineage. The underlying reproductive isolation mechanisms may involve a combination of intrinsic and extrinsic genetic barriers, where relative contributions may differ between the two contexts.

### 1-Genome scans in hybrid zones

Our SNP panel allowed us to define five discrete panmictic genetic clusters, two in the Atlantic Ocean, two in the Mediterranean Sea, and one in the Black Sea, which cannot be morphologically distinguished with reliability so far (*i.e.* cryptic lineages). The average genetic differentiation between clusters and its associated variance were strong (0.09 ± 0.02 < *F*_*ST*_ ± sd < 0.26 ± 0.04). In this context, identifying outlier loci is a complex task with problems of false positives (Lotterhos and Whitlock 2014, 2015). It is also increasingly being acknowledged that a signal of local adaptation is not easily captured without a broad sampling of the genome in a standard infra-specific low-linkage disequilibrium context (Hoban et al. 2016). Finding easily nine well-supported outlier loci with a limited number of loci suggests strong variance in differentiation levels associated with the existence of cryptic genetic barriers involving many selected loci (Bierne et al. 2011). Extensive linkage disequilibrium is also maintained in this complex of semi-isolated genetic backgrounds when compared to a standard infra-specific context. However, this is not expected to be a rare situation as genomic studies have provided accumulating evidence that semi-permeable barriers to gene flow are widespread (Roux et al. 2016) and affect a substantial proportion of genomes (Harrison and Larson 2016).

In this study, six of the nine outliers showed a signature of genetic parallelism (Fig. 2, Fig. SI3). These six outliers not only proved to be the most differentiated loci using a pairwise comparison between Northern and Southern lineages in the Atlantic Ocean and between the lagoon and marine ecotypes in the Mediterranean Sea (Fig. 6), but they also displayed a genetic structure that is not compatible with the sample covariance matrix inferred with the full set of loci (FLK test, Supporting Information SI6) or with simulations under the best-supported demographic model inferred from transcriptome data (Roux et al. 2016). Finally, these six outliers (3.15% of the SNPs analyzed) proved to map to the same chromosome (4.5% of the genome). These six loci mapped to the same chromosome in available fish genomes, although the order on the chromosome was not so well conserved in distantly related species. This result suggests the existence of a large genomic island of differentiation as already reported in other fishes (e.g. sticklebacks: Jones et al. 2012, cod: Hemmer-Hansen et al. 2013, Berg et al. 2017, seabass: Duranton et al. 2018), and possibly involving a chromosomal inversion or another form of recombination suppression. This chromosome-wide island of differentiation also explains why we detected the signal of parallel outlying differentiation with a moderate number of loci analyzed. The chromosome could have been easily missed with a few microsatellites but our SNP panel was largely enough to have a few SNPs on the chromosome. These preliminary results call for genome sequence analysis in order to better characterize the genomic island and its structural variation, and to investigate the genome localization of additional barrier loci with a smaller chromosomal footprint.

### 2-Parapatrically distributed semi-isolated cryptic lineages in the Atlantic Ocean

In the Atlantic Ocean we confirmed the existence of two well-differentiated genetic clusters previously identified by Woodall et al. (2015) with five microsatellites and two mitochondrial genes. A genetic homogeneity was observed over large distances within each lineage (Fig. 1), contrasting with a very strong and abrupt genetic differentiation between them. Inter-lineage genetic divergence was not only captured by the most differentiated SNP that were nearly fixed between the two lineages (SNP 130 in SI4, *F*_*ST*_ = 0.9), but also, by many other SNPs (SI3B), suggesting isolating mechanisms and loci with genome-wide effects. This mechanism is also suggested by the discovery of two coexisting lineages in the same lagoon in South West of France without early generation hybrids sampled (Hossegor lagoon, site 6 in Fig. 1). Such zone of co-existence in sympatry contrasting with genetic homogeneity at a large spatial scale within each lineage suggests strong reproductive isolation (Jiggins and Mallet 2000), involving either pre-zygotic isolation or strong selection against hybrids at early life stages.

Recent migration of the two diverged genetic backgrounds in this lagoon does not easily explain the co-existence of the two lineages with no hybridization. In addition to the low probability of sampling the first-generation migrants without any genomic modifications, local introgression of the North-Atlantic seahorse in this lagoon (Fig. 3) attests hybridization occurred and that a mechanism of isolation must maintain the divergence. The decreasing proportion of south lineage ancestry from Hossegor to the English Channel also provides indirect evidence for asymmetric introgression (Fig. 2, SI3A). In addition, an interesting pattern was observed at genomic island outlier loci in Hossegor (site 6) that further support partial reproductive isolation. The North Atlantic lineage mostly carries the southern alleles at genomic island loci (pale pink dots in Fig. 4, see also SI3B). To produce such North Atlantic-type individuals with a South-Atlantic island, recombination between parental backgrounds is required, suggesting hybridization occurred in this case. In the genomic cline analyses (Fig. 7) the geographic information was included to contrast the results obtained with or without the Hossegor sample. The objective of this analysis was to infer the existence of a local discordance in this site. The swamping of North Atlantic fishes by the southern allele at the genomic island suggests epistatic or coupling interactions among loci implied in reproductive isolation. Endogenous post-zygotic selection against hybrids does not result in a stable polymorphism in a single isolated population, but instead in transient polymorphisms. Also known as bistable variants, the maintenance of underdominant or epistatically interacting genetic incompatibilities requires a migration-selection balance in a spatially subdivided population (Barton and Turelli 2011), or frequency-dependent selection (Barton and de Cara 2009). The system may have fixed one state of a bistable variant in the Hossegor lagoon. Alternatively, the southern allele may confer a selective advantage in this lagoon whatever the genetic background. However the latter hypothesis does not seem coherent as the northern allele is always in higher frequency in lagoon-like habitats elsewhere, though we may have missed more subtle ecological parameters that would have driven this pattern.

Overall, the Atlantic contact zone possesses all the characteristics of standard clinal hybrid zones that follow the tension zone model (Barton and Hewitt 1985), *i.e.* a secondary contact zone maintained by a balance between migration and intrinsic reproductive isolation. Exogenous selection may also contribute, although the two Atlantic lineages both inhabit indifferently lagoon and sea habitats (Fig. 1), and co-exist in syntopy in Hossegor, suggesting a limited role of ecological contrast in their genetics. In any case, only strong intrinsic reproductive isolation can guarantee a genome-wide barrier to gene flow explaining the co-existence of the two lineages in sympatry in the Hossegor lagoon.

### 3-Sea and lagoon ecotypes in the Mediterranean Sea

Contrasting with the Atlantic hybrid zone, the two cryptic lineages identified in the Mediterranean Sea were associated with lagoon/sea ecosystem variation. Our broad genomic and spatial sampling revealed two crucial observations. First, while the marine lineage was surprisingly homogeneous over the whole Mediterranean Sea, from Greece to Spain (*F*_*IT*_ = 0.0078 n.s.), lagoon-like samples, especially the Thau lagoon, showed a strong and genome-wide genetic differentiation from them. Samples from two lagoons (Thau in France and Bizerte in Tunisia, sites 16 and 20 in Fig. 1) were sufficient to reveal an association with the environment that was previously unseen, the Thau lagoon being the only sample from Western Mediterranean basin in Woodall et al. (2015). Second, fixed differences between lagoon and marine samples were observed, although they were sampled only few kilometers apart (e.g. sites 14-16 in Fig. 2, 4). A single but important seahorse sampled on the seaside of the Thau lagoon (site 15 in Fig. 1), plus seven others sampled on the seaside of another lagoon of the region (site 14 in Fig. 1) proved to belong to the marine genetic cluster, without any sign of introgression. Likewise, no evidence of introgression was observed in the Thau sample (Fig. 3). Once again, despite genetic homogeneity over large area, such strong and abrupt genetic differentiation suggests partial reproductive isolation between these two lineages. In this case there are obvious ecological drivers, *i.e.* habitat specialization. Indeed, the Bizerte lagoon (site 20 in Fig. 1), which is ecologically similar to the Thau lagoon (Sakka Hlaili et al. 2008), has a population genetically similar to marine Mediterranean samples at most loci and only share the genetic composition of the Thau lagoon at the genomic island (Fig. 4 and SI3B). In addition, a subsample of the Halkida population (Greece, site 24B in Fig. 4) was composed of five individuals heterozygous at the genomic island. The environmental parameters at this location are hypothesized to be more lagoon-like, being a secluded bay beyond the northern end of the Euipus Strait. This sample only provides evidence that the genomic island polymorphism and the mosaic spatial structure extends to the eastern basin without really providing further clues about the role of the environment.

Our results from the Mediterranean Sea (*i.e.* lagoon/marine system) resemble those obtained from the emblematic three-spined sticklebacks marine/freshwater system (Jones et al. 2012), or more recently from the marine-migratory/freshwater-resident lampreys (Rougemont et al. 2016) and coastal/marine ecotypes in European anchovies (Le Moan et al. 2016). When a shared divergent genomic island is observed –*i.e.* genetic parallelism, as here in long-snouted seahorse among the Mediterranean lagoons, or in the examples cited above, there could be three possible interpretations: (i) parallel gene reuse from a shared ancestral polymorphism present in the marine supposedly ancestral population (Jones et al. 2012), (ii) the spread of a locally adapted allele (*i.e.* the ‘transporter hypothesis’; Schluter and Conte 2009) or (iii) secondary contact followed by spatial reassortment of the divergent lineages and extensive introgression swamping such that only selected loci and their chromosomal neighborhood retain the history of adaptation (Bierne et al. 2013). The three scenarios are difficult to discriminate as they converge toward a similar pattern (Johannesson et al. 2010, Bierne et al. 2013, Welch and Jiggins 2014). Here, as for the lampreys (Rougemont et al. 2016), the Thau lagoon provides a possible support for the secondary contact model because the differentiation, although stronger at the genomic island, is genome-wide. The Bizerte lagoon however can either be interpreted as a marine lineage introgressed by the lagoon allele at the genomic island, or as a lagoon lineage (defined by adaptive/speciation genes) massively introgressed by neutral marine alleles. Incorporating heterogeneous migration rates in demographic inference methods allowed Le Moan et al. (2016), Rougeux et al. (2016) and Rougemont et al. (2016) to identify the signal of a secondary contact history carried by islands of differentiation in lampreys, white fishes and anchovies. Unfortunately our 286-SNPs dataset does not allow performing such historical demographic reconstruction. Nonetheless the TREEMIX analysis reveals that episodes of secondary admixtures strongly improve the fit to the sample covariance matrix, but adaptive introgression or massive introgression swamping can both explain them. Anyhow, demographic reconstruction does not completely refute the ‘transporter hypothesis’ which stipulates lagoon alleles spread from lagoon to lagoon (or freshwater allele from river to river) and is a scenario that produces a very similar genomic pattern of differentiation to the one produced by a standard secondary contact (Bierne et al. 2013, Rougemont et al. 2016). In the case of the seahorse complex, however, we made the new observation that genetic parallelism is observed with the Atlantic populations where the structure is geographic and independent of the lagoon-sea habitats, which offers complementary arguments to the debate.

### 4-Genetic parallelism in two different spatial/ecological contexts

The most astonishing result of our genetic analysis was genetic parallelism between the Mediterranean lagoon ecotype and the north Atlantic lineage at a large genomic island. Parallel evolution implies distinct but repetitive ecological characteristics (e.g. Butlin et al. 2014). However, this claim comes from an inductive reasoning, validated by many observations, which a single counter-example can yet alone refute. In the present study, we found that the genomic island was associated with the sea-lagoon ecological contrast in the Mediterranean Sea, while there was no such genetic differentiation between lagoon and sea samples in the Atlantic Ocean. The North and South lineages inhabit indifferently lagoons and seas, so that what seemed obvious in the Mediterranean Sea regarding the divergence of the two lineages, *i.e.* habitat specialization, was not observed along the Atlantic Ocean where the divergence seems uncorrelated to the ecological contrasts that explain the two Mediterranean ecotypes. Besides, abiotic parameters such as temperature or salinity, did not allow bounding the North Atlantic and Mediterranean lagoon lineages genetic parallelism. Although we may have missed putative ecological drivers of such genetic parallelism, parallel gene reuse driven by ecological convergence seems here unlikely. Shared ancestral polymorphism sieved by adaptation in a patchy environment (Bierne et al. 2013) and incipient speciation (Guerrero and Hahn 2017) would better explain our data. The association with habitat in the Mediterranean Sea and with space in the Atlantic Ocean could be explained by a secondary evolution of locally adapted genes within the genomic island in the Mediterranean Sea, benefiting from the barrier to gene flow imposed by intrinsic selection (divergence hitchhiking; Via 2012, Yeaman 2013). Alternatively, the genomic island could have coupled with local adaptation polymorphisms localized elsewhere in the genome in the Mediterranean Sea (Bierne et al. 2011), while it would have been trapped by a barrier to dispersal in the Atlantic Ocean (Barton 1979, Barton and Hewitt 1985). Without further data and the true genomic position of loci in the seahorse genome, it is difficult to disentangle the two hypotheses.

One locus (SNP 87) localized in the same chromosome as parallel SNPs (Fig. 5) is differentiated between the marine and lagoon ecotypes in the Mediterranean Sea, but is not differentiated between the northern and southern lineages in the Atlantic Ocean. At first sight, it could be interpreted as evidence for a possible secondary local sweep in Mediterranean lagoons. However, according to the gene order inferred from the closest species (Gulf pipefish and Tiger tail seahorse) along with a barrier to gene flow less effective in the Atlantic (see SNPs 34 and 212 in Fig. 4 and 5B), it could also be interpreted as being localized in the island “shoulders” (*i.e.* loci in the vicinity of the regions harboring local adaptation and/or reproductive isolation loci; Gagnaire et al. 2015, Le Moan et al. 2016) in which a stronger introgression rate has erased the differentiation faster in the Atlantic than in the Mediterranean, in favor of the alternative interpretation. Importantly, whatever the explanation-divergence hitchhiking or coupling-it requires invoking interaction between intrinsic and ecological selection and not ecological selection alone (Bierne et al. 2011, Kulmuni and Westram 2017). Furthermore, intrinsic selection has most probably evolved first in this system as no genetic parallelism was observed in outliers discriminating lagoon to marine ecotypes, which would contradict the predominant view that ecological selection is necessarily the initial catalyzer of the chain of accumulation of barriers in ecological speciation.

## Conclusions

Analyzing the population genetics of the long-snouted seahorse *Hippocampus guttulatus* revealed a complex of panmictic genetic backgrounds subdivided by sharp semi-permeable hybrid zones. This is now standard observation in marine species (Knowlton 1993, Pante et al. 2015, Sheets et al. 2018) where morphological stasis might be more widespread than in terrestrial species. The subdivision of species by hybrid zones is a long-lasting observation in the terrestrial realm (Hewitt 1989) but arguably more readily detected by morphological differences. We also easily found outlier loci despite a moderate number of loci analyzed, and the clustering of these outlier loci in a single genomic region. This also tends to become a standard observation of the recent hybrid zone literature (e.g. in stickelbacks, Jones et al. 2012; jaera, Ribardière et al. 2017; cod, Hemmer-Hansen et al. 2013).

However, we also made two additional observations that are less commonly reported and deserve a broader interest outside the study of seahorse themselves, although are important to further inform captive breeding and *in-situ* management decisions. First, we found the two, usually opposed, standard spatial structures of the hybrid zone literature, namely clinal and mosaic hybrid zones, in the same study system. This result calls for further investigations with lab and fieldwork in order to better understand the mechanisms of reproductive isolation at play and their genetic architecture. Secondly, we found a parallel pattern of differentiation at the genomic island in the two spatial/ecological contexts. Although this result will also need to be substantiated by follow-up genomic studies, it nonetheless reveals that the hallmark of ecologically driven adaptive divergence can be observed in absence of obvious ecological convergence. We argue that alternative scenarios involving secondary introgression swamping and intrinsic isolation should be more seriously considered as valid alternatives and the seahorse complex could become an interesting flagship system in the debate.

## Acknowledgments

We are grateful to Lucas Beranger, Michel Cantou, Philippe Lenfant, Pablo Liger, CPIE Bassin de Thau, Patrick Lelong, Francesco Di Liello and Stéphane Auffret for their help in providing *H. guttulatus* fin-clippings and to Fabienne Moreau for the BeadXpress experiment. Many thanks to Nicolas Duforet-Frebourg and Christelle Fraïsse for computational advice. This work was funded by a Languedoc-Roussillon Region “Chercheur(se)s d’avenir” grant to NB (Connect7 project), by a LabEx CeMEB postdoctoral fellowship to FR and by Chocolaterie Guylian and a Natural Environment Research Council Industrial Case studentship (NER/S/C/2005/13461) to LW.

## Data archival location

SNPs data have been deposited at DRYAD. DOI: 10.5061/dryad.mq122fv.

